# Identifying Migrant Origins Using Genetics, Isotopes, and Habitat Suitability

**DOI:** 10.1101/085456

**Authors:** Kristen C. Ruegg, Eric C. Anderson, Ryan J. Harrigan, Kristina L. Paxton, Jeff Kelly, Frank Moore, Thomas B. Smith

## Abstract

1. Identifying migratory connections across the annual cycle is important for studies of migrant ecology, evolution, and conservation. While recent studies have demonstrated the utility of high-resolution SNP-based genetic markers for identifying population-specific migratory patterns, the accuracy of this approach relative to other intrinsic tagging techniques has not yet been assessed.
2. Here, using a straightforward application of Bayes' Rule, we develop a method for combining inferences from high-resolution genetic markers, stable isotopes, and habitat suitability models, to spatially infer the breeding origin of migrants captured anywhere along their migratory pathway. Using leave-one-out cross validation, we compare the accuracy of this combined approach with the accuracy attained using each source of data independently.
3. Our results indicate that when each method is considered in isolation, the accuracy of genetic assignments far exceeded that of assignments based on stable isotopes or habitat suitability models. However, our joint assignment method consistently resulted in small, but informative increases in accuracy and did help to correct misassignments based on genetic data alone. We demonstrate the utility of the combined method by identifying previously undetectable patterns in the timing of migration in a North American migratory songbird, the Wilson's warbler.
4. Overall, our results support the idea that while genetic data provides the most accurate method for tracking animals using intrinsic markers when each method is considered independently, there is value in combining all three methods. The resulting methods are provided as part of a new computationally-efficient R-package, GIAIH, allowing broad application of our statistical framework to other migratory animal systems.

## 1. introduction

The ecology and evolution of animals that undergo annual seasonal migration is shaped by events encountered across the entire annual cycle (Sillett *et al*., 2000; Webster *et al*., 2002, 2005). It is now well established that habitat conditions during migratory or wintering phases can have significant carry-over effects on breeding ground productivity (Marra *et al*., 1998; Sillett *et al*., 2000; Norris & Taylor, 2006). As a result, understanding patterns of migratory connectivity, or the geographic links between breeding, wintering, and stopover sites for a population over the course of an annual cycle, is a critical first step towards studies of migrant ecology, evolution, and conservation.

Efforts to identify the strength of migratory connections have relied on a variety of methods for tracking animal movements (Marra *et al*., 1998; Bonfil *et al*., 2005; Smith *et al*., 2005; Stutchbury *et al*., 2009). Extrinsic devices such as satellite transmitters and geo-locators have increased our knowledge of the movement patterns of individuals of a particular species (Stutchbury *et al*., 2009), but remain impractical for many large-scale (1000’s of individuals) applications due to cost and weight restrictions, and the need to re-capture many individuals to collect the data (Arlt *et al*., 2013; Bridge *et al*., 2013). An attractive alternative is the use of genetic and isotopic markers, that capture information contained within the tissue of an animal, to pinpoint an individual’s population of origin. These methods have broad appeal because they are minimally invasive, cost-effective when applied at scale, and do not require recapture of individuals (Rubenstein *et al*., 2002; Kelly *et al*., 2005; Rundel *et al*., 2013). Furthermore, intrinsic methods make it possible to trace the origins of animals that have died from both natural and anthropogenic causes (i.e., poaching, collisions, and disease), because genetic and isotopic samples can be collected from carcasses.

While some genetic approaches have been limited by a lack of resolution, advances in genome-wide sequencing have resulted in new technologies that can be applied to genetic tagging of wild populations (Allendorf *et al*., 2010; Metzker, 2010; Davey *et al*., 2011). Even in species with high rates of dispersal, such as birds, fish, and mammals it has been found that a small number (*n* <100) of single nucleotide polymorphisms (SNPs) found within, or linked to, genes under selection can be targeted to reveal population structure at spatial scales that are critical to regional conservation planning (Nielsen *et al*., 2009; Hess *et al*., 2011; Nielsen *et al*., 2012; Ruegg *et al*., 2014). For example, Ruegg et al (2014) found that 96 high-resolution SNPs could be used to identify six genetically distinct populations of a migratory songbird, the Wilson’s warbler (*Cardellina pusilla*), whereas previous single-marker techniques found support for only two groups (Kimura *et al*., 2002). Furthermore, SNP assays that isolate short fragments of DNA specific to the taxa of interest make it possible to rapidly screen DNA from a variety of tissue types (i.e., bird feathers, fin clips, animal hair) that can be collected using non-invasive sampling techniques, making this method an attractive choice for conservation genomic studies (Ruegg *et al*., 2014; Kraus *et al*., 2015).

Despite their appeal, several questions remain as to how genetic tools compare in accuracy with other intrinsic marking methods. In addition, there are clear examples where genetic markers alone fall short of resolving populations across all or large parts of a species geographic range (Gagnaire *et al*., 2015; Toews *et al*., 2015). In such situations, the inclusion of additional sources of non-genetic information, such as isotopic data and/or habitat suitability models, can increase the resolution of genetic markers (Kelly *et al*., 2005; Rundel *et al*., 2013; Pekarsky *et al*., 2015). For example, Rundel et al. (2013) showed that genetic and isotopic information can be combined to increase the assignment accuracy of individuals (birds) to their population of origin; however this method was not designed to deal with disjunct patterns of genetic variation, like that observed in the Wilson’s warbler (Ruegg *et al*., 2014). More recently, Pekarsky et al. (2015) used habitat suitability models to refine their isotopic-based estimates of population assignment, but did not establish whether the use of habitat suitability as a prior actually led to improved estimates of the geographic origin of individuals.

Here we combine genetic, stable isotope, and habitat suitability data into a joint assignment procedure that infers the breeding origins of individuals collected anywhere along their migratory trajectory with greater resolution than can be attained using each method individually. We refine previously developed R-code (R Core Team, 2016) for performing each type of assignment alone (Anderson *et al*., 2008; Bridge *et al*., 2013; Vander Zanden *et al*., 2014), making it computationally feasible to combine assignments and perform statistically rigorous leave-one-out cross validation. To assess the overall accuracy of our method in comparison to existing approaches we examine the results from each type of data—genetics, stable isotopes, and habitat suitability—individually as well as jointly, in order to determine: (1) contributions of each data type to the accuracy of joint assignments, (2) insights that might be gathered from the combined approach, and (3) recommendations for future studies of migratory connectivity based on our results.

We assess the accuracy of each method using data from a long-distance migratory bird, the Wilson’s warbler (*Cardellina pusilla*), a particularly appropriate model for testing the efficacy of our approach because previous population-genetic/connectivity studies on this species provide a solid basis for comparison between methods (Kimura *et al*., 2002; Clegg *et al*., 2003; Paxton *et al*., 2007; Irwin *et al*., 2011; Paxton *et al*., 2013; Rundel *et al*., 2013). We start by deriving posterior probability rasters (or other scaled “scores”) for each method individually (genetics, stable isotopes, and habitat suitability) and then combine those into a joint assignment probability. We then evaluate the gains in accuracy achieved by each method individually as well as by combining multiple data sources using leave-one-out cross-validation with a reference set of Wilson’s warblers sampled from known locations during the breeding season. The combined approach is then applied to the assignment of migratory birds of unknown origin at a stopover site in Cibola, AZ during spring migration. Our methods have been implemented in the new R-package, GAIAH (**G**enetic **A**ssignment using **I**sotopes **A**nd **H**abitat suitability), which also includes all the data and scripts required to replicate our results in this paper. It is available on GitHub (https://github.com/eriqande/gaiah).

## 2. Methods

### 2.1 Sampling

Sampling of Wilson’s warblers is detailed in Ruegg *et al*. (2014). Briefly, collection of 357 feathers from 30 locations (average of 12 individuals/site; range: 2-25) across the breeding range was made possible through a large collaborative effort with bird banding stations both within and outside of the Monitoring Avian Productivity and Survivorship (MAPS) and the Landbird Monitoring of North America (LaMNA) networks. These samples became the foundation of the subsequent genetic and isotopic analysis of known-origin samples. Breeding samples were collected and categorized into groups based on collection date (June 1 to July 31), signs of breeding (presence/size of a cloacal protuberance), and life history timetables for the Wilson’s warbler. To illustrate the efficacy of the combined approach to the assessment of migratory stopover site use-through-time, 686 migrant samples were also collected from Cibola, AZ (31°18'N, 114°41'W), using consistent-effort, daily, passive mist-netting from March 22 to May 24, in both 2008 and 2009.

### 2.2 Genetics

The application of genetic markers to the assignment of Wilson’s warblers is detailed in Ruegg *et al*. (2014). Briefly, genetic samples, consisting of the proximal end of one rectrix (breeding samples) and whole blood samples (migratory samples) were purified using a Qiagen DNeasy Blood and Tissue Kit and quantified using a NanoDrop™ Spectrophotometer (Thermo Scientific, Inc) (Smith *et al*. 2003). Genotyping was done using a panel of 96 high-resolution SNP markers ascertained from a genome-wide survey of genetic variation in the species using restriction-associated-digest, paired-end (RAD-PE) sequencing. Analyses using structure (Pritchard *et al*., 2000; Falush *et al*., 2003) and geneland (Guillot *et al*., 2005) indicated G = 6 spatially-distinct population groups that could be reliably distinguished using the 96 SNPs. The spatial extent of these six groups identified by GENELAND was overlaid, and then clipped, by the known range of Wilson’s warbler [downloaded from NatureServe, www.natureserve.org, (Ridgely *et al*., 2005)], to create a map of the occurrence of different population groups across the breeding range (See Figure 1, Ruegg *et al*. 2014). Breeding birds collected from locales within each population group were included as reference samples for genetic population assignment to assign posterior probabilities across the 6 genetic groups using the program gsi_sim (Anderson *et al*., 2008). For each bird, i, this program returns an estimate of the posterior probability,

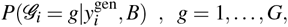

That *i*’s genetic group (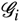) is equal to *g* (*g* denotes one of the six genetic groups), given *i*’s genotype 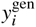, and the reference or baseline samples, *B*.

These posterior probabilities refer to group membership of each bird; however, to combine these inferences with stable isotope data requires first converting group membership posteriors into posterior probabilities of spatial location. As the 96 SNPs provide only limited ability to resolve local origin of birds *within* each genetic group’s geographic area, we converted the genetic posteriors for the *i*^th^ bird to a grid, 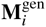, of spatially-explicit posterior probabilities by distributing the posterior probability 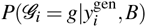, *B*) uniformly across the raster grid representing the spatial range of group *g*, and ensuring that these are appropriately normalized to sum to one across the whole breeding range. Namely, within group *g*’s range, posterior probability is distributed according to

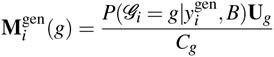
where **U**_*g*_ is a matrix (raster) containing 1’s in cells within genetic group *g*’s region and 0’s elsewhere, and *C_g_* is the number of raster cells within group g’s region (*i.e.*, the sum of the elements of U_*g*_). In most cases within the reference dataset indivdiuals were assigned with high probablity to one genetic group and lower probabity to other groups (see Figure 1, Ruegg *et al* 2014). To represent the uncertainty in the genetic assignments to different groups (which in most cases was very small, see Supporting Information Document 1, Figure 1) we spread the posterior probability of assignment to each group across the geographic ranges of all the groups to which the bird was assigned and then weighted the values according to their likelihood so that the combined areas summed to 1.

Subsequently, the posterior probability of spatial assignment of birds based on genetics and assuming a uniform prior across space is

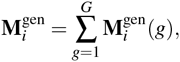

Whose elements clearly sum to unity. 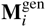 is a raster of the same extent and resolution as 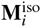 (see next section).

### 2.3 Stable Isotopes

Predictable continental patterns of stable hydrogen isotopes in rainfall (*δ*^2^H_*p*_) are highly correlated with stable hydrogen isotopes of animal tissues (*δ*^2^H_*f*_), allowing for inferences about the origin of where tissues were grown (Hobson & Wassenaar, 2008). When the breeding and molting locations are the same, as is the case for Wilson’s warblers, then stable isotope values provide an assessment of the breeding origin of a bird that is independent of a genetic assessment. We expressed all isotope ratios in standard delta notation (*δ*^2^H) where

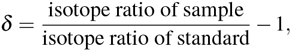

With ratios shown as parts per thousand (‰) deviation from Vienna Standard Mean Ocean Water for hydrogen. Prior to analysis, all feathers (breeding and migratory samples) were cleaned with dilute detergent followed by a 2:1 chloroform:methanol solution (Paritte & Kelly, 2009). For *δ*^2^H analyses, a 0.1-0.2 mg piece of feather was packed into a silver capsule and loaded into an auto-sampling tray. Isotope ratio measurements were performed at the University of Oklahoma with a ThermoFinnigan Delta V isotope ratio mass spectrometer connected to an elemental analyzer (H analyses: TC/EA, Thermo-Finnigan). To control for exchangeable hydrogen, hydrogen isotope ratios were normalized according to Wassenaar & Hobson (2003), using established keratin standards: chicken feathers (-147.4‰), cow (*Bos taurus*) hooves (-187‰), and bowhead whale (*Balaena mysticetus*) baleen (-108‰). For additional details on our analysis methods, see Kelly et al. (2009) and Paritte & Kelly (2009).

We created an isoscape of *δ*^2^H_*p*_ ratios and its associated variance using IsoMAP (Job Key 54152), an online resource to generate region-and time-specific isoscapes for geographic assignments (Bowen *et al*. 2014, www.isomap.org). In IsoMAP, a geospatial isoscape was generated using precipitation data from 120 stations collected during the time period of 1960-2009, and included CRU-derived climatic variables such as elevation, precipitation, and minimum precipitation in the model (Mitchell & Jones 2005; similar to Hobson *et al*. 2012c). Using a parametric bootstrapping approach we converted the isoscape of *δ*^2^H_*p*_ values to an isoscape of *δ*^2^Hf values based on the relationship between *δ*^2^H_*p*_ and 5^2^H*f* collected from the 357 known-origin, breeding Wilson’s warblers sampled across the breeding range (Appendix 1). We then computed the posterior probability of breeding origin given a feather *δ*^2^H*f* ratio and three sources of variance following the methods used in Vander Zanden et al. (2014). The three sources of variance included variance associated with; 1) the original *δ*^2^H_*p*_ isoscape generated in IsoMAP, 2) the rescaled precipitation to feather *δ*^2^H isoscape, and 3) individual variation in *δ*^2^H*f* among birds sampled at the same breeding location.

Specifically, this approach assumes that 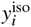, the isotope ratio measured in the feather of the *i*^th^ bird, is a normal random variable with mean and variance determined by its location in space. If these means and variances, denoted by the matrices **T**^(μ)^ and **T**^(σ^2^)^, respectively, were known across a regular grid of possible origin locations, then it would be straightforward to compute the posterior probability of breeding location. Namely, assuming a uniform prior on spatial origin, the posterior that bird i originated from the (*r, c*)^th^ cell in the grid is proportional to the density of observing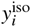given it was drawn from a normal density with mean 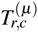 and variance 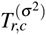 (the (*r,c*)^th^ elements of **T**^(μ)^ and **T**^(σ^2^)^,respectively).

It should be noted that**T**^(μ)^ and **T**^(σ^2^)^ are not known, so we follow Vander Zanden *et al*. (2014) by using 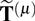 and 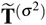 in their place, computed as
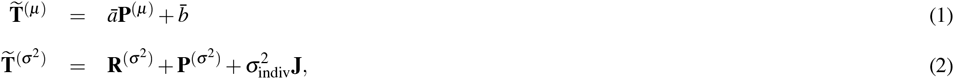

Where **P**^(μ)^ and **P**^(σ^2^)^ are the predictions and associated variances, respectively, for the precipitation isotope ratios from IsoMAP made on the same grid as 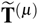 and 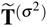is a matrix of 1’s of the same dimension as 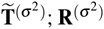 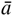, and 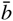, are determined by a parametric bootstrapping approach described in Appendix 1; and 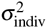is a term accounting for individual variation in isotope ratios which was set to the square of the mean of the standard deviation of 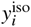amongst birds sampled at common locations across the breeding range during the breeding season.

Posterior probabilities derived in this manner for bird *i* are a set of values over a spatial grid denoted by the matrix, 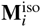

### 2.4 Habitat Suitability

In order for subspecies ranges to be further classified in terms of their utilization by breeding individuals, habitat suitability models were constructed for Wilson’s warblers across their breeding range. This was performed by identifying the geographical locations of Wilson’s warblers during the breeding season, and then modeling the relationship of these occurrences to the environmental conditions at those sites. This model was then used to predict the probability of occurrence of individuals across the species range.

Locations of individuals were extracted from the eBird website (www.ebird.org). To ensure that we used only those birds that were breeding (or at least were located on breeding grounds), we filtered the nearly 450,000 records of Wilson’s warblers available to include only those sighted on or after June 10, but before August 1, of any given calender year. To avoid redundant observations at the same site artificially influencing models, we used each unique geographic location only once. The final dataset contained 9,984 unique locations of Wilson’s warblers on breeding grounds.

We used the machine learning algorithm MaxEnt (Phillips *et al*., 2006) to model Wilson’s warbler distributions. As this method relies only on presence data (*i.e.*, it does not require records of Wilson’s warbler *absence*), it is particularly well-suited for the use of eBird occurrence records to capture complex biological responses to environment (Elith *et al*., 2011), and has performed well in previous statistical comparisons with other species distribution modeling techniques (Harrigan *et al*., 2014). Environmental variables were chosen to best reflect unique predictive ability, and variables were removed from final analyses when a Pearson’s correlation coefficient was >0.7 among the variable and any other predictor. We ran 3 replicates of a MaxEnt model using the Wilson’s warbler records as a response variable and 14 climate and landscape variables as predictors. These included 8 (among 19 available) bio-climatic layers (Bio 1,2,4-6,12,15,19, downloaded from www.worldclim.org) as well as variables representing landcover, tree cover, elevation, and vegetation characteristics and heterogeneity [Normalized Difference Vegetation Index (NDVI) mean and standard deviation] downloaded from the Global Land Cover Facility (www.landcover.org). Default parameters were used for all MaxEnt runs except for the following: clamping turned off, a maximum of 100,000 background points selected, and 20% of data withheld for subsequent testing. In addition, both predictor response curves, as well as jackknifing to assess variable importance, were created as part of the output.

Once final models were evaluated and found to converge on similar response curves and probability estimates, the mean output ascii file was exported to ArcGIS and was clipped to the extent of the breeding range of Wilson’s warbler, as determined using digital range maps provided by NatureServe (www.natureserve.org, (Ridgely *et al*., 2005)). These final probability estimates were then downsampled in resolution to yield a raster **M**^hab^ with the same extent and resolution as 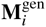 and 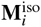.

### 2.5 Combining data types

The above approaches yield 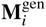and 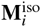 for the *i*^th^ bird, as well as a prior probability of occurrence of a Wilson’s warbler in any location, 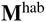. Treated as matrices of probabilities, these rasters can be combined using Bayes’ rule to obtain a combined estimate of probability of spatial origin for each bird. In order to explore weighting the different sources of information (genetics, stable isotopes, and habitat) differently, we considered incorporating the parameter 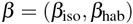 that determines the relative amount of weight given to the stable isotope and habitat data, and hence compute the combined probability as
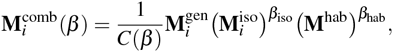

Where (**M**)*^β^* denotes the matrix **M** with every element raised to the power *β*. *C*(*β*) is a normalizing constant that ensures that the elements of 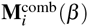 sum to one. It can be easily found for any 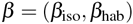 by simply summing over all the elements of the unnormalized matrix.

Among the different genetic groups of birds, the relative utility of isotopic data versus habitat suitability may vary. In theory, one could leverage this to advantage by choosing different values of *β*_iso_ and *β*_hab_ for birds that are assigned with genetics to particular genetic groups. However, doing so accurately would require sampling of reference birds proportionally with respect to their true density on the landscape, which may not be the case with the genetic samples used herein, which were obtained opportunistically from bird banding stations. Accordingly, while we explored a range of values of (*β*_iso_, *β*_hab_), for all results in the paper we used values of *β*_iso_ = *β*_hab_ = 1.

### 2.6 Assessment of Accuracy

To assess the extent to which the inclusion of stable isotope and habitat data improve the spatial localization of birds on the breeding grounds we test how well birds of known breeding location (those in the reference data set) can be inferred using genetics alone, stable isotopes alone, habitat suitability alone, or a combination of each. When doing so with the reference birds, we use a leave-one-out procedure, removing individual *i* from the reference data set when computing 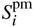 and 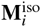.

To assess accuracy we develop an easily interpreted metric, 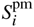, the posterior mean of great circle distance between the true location of *i* and inferred locations. This is found by averaging the distance between *i*’s true location and the center of every cell in the breeding range, weighted by **M**_*i*_. In order to assess the accuracy of genetics, stable isotopes, or habitat used separately or together, we compute 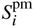 with 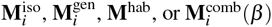, respectively.

### 2.7 Assignment of Unknown Migratory Birds-Example Dataset

To illustrate the efficacy of the combined approach we applied the combined approach to a set of migratory birds of unknown origin captured at a stopover site in Cibola, AZ during spring migration. For each bird, we determined the combined probability of origin using equation 3, creating a matrix of probability assignments, 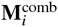, for each migrant. We then calculated the posterior mean migration distance remaining from the stopover site by averaging the distance between the stopover site and the center of every cell in the breeding range, weighted by 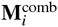.

## 3 Results

### 3.1 Genetics

As previously reported (Ruegg *et al*., 2014), the *maximum-a-posteriori* self-assignments of the reference birds using genetic data showed that a high fraction (88.2%) of birds are correctly assigned to their true region of origin. The rate of correct assignment varied across regions, with the lowest rate of correct assignment occurring in Coastal California (78%), followed by Rocky Mountain (81%), Pacific Northwest (85%), California Sierra (86%), Alaska to Alberta (95%), and Eastern North America (100%). Of the birds that are assigned correctly, their average maximum posterior is 96% and when birds are incorrectly assigned to their true region of origin, the average maximum posterior is 74%. This confirms that when genetic assignments are incorrect, isotopes and suitability have the potential to increase posteriors, and thereby improve localizations in joint assignments.

### 3.2 Stable Isotopes

Across the Wilson’s warbler breeding range, *δ*^2^H_*p*_ values range from -14.3‰ to -190.6‰ with uncertainty across the *δ*^2^H^*p*^ isoscape varying (SD: 8.53 to 21.64) as a result of the uneven distribution of precipitation stations used to model *δ*^2^H_*p*_ ratios. Regions with the greatest uncertainty in *δ*^2^H_*p*_ values included coastal Alaska, Northwest Territories, and Newfoundland. The rescaling equation used to convert the *δ*^2^H_*p*_ isoscape to a *δ*^2^H_*f*_ isoscape was *δ*^2^H_*f*_ = 0.74(*δ*^2^H_*p*_) -38.01; that is, in Equation 1, a = 0.74 and b = -38.01. The standard deviation associated with the rescaled precipitation to feather *δ*^2^H isoscape, **R**^(σ)^, ranged from 0.61 to 6.56, while the mean, within-site standard deviation of *δ*^2^H_*f*_ from the 357 Wilson’s warblers of known breeding origin sampled across 30 locations was **σ**_indiv_ = 12.49 (Supplement Table 1). The total variability, 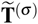, including the three sources of uncertainty, contained standard deviation values ranging between 15.15 and 24.99, depending on the geographic region of assignment.

### 3.3 Habitat Suitability

Habitat suitability model runs yielded accurate species probability of occurrence maps according to several criteria. First, the mean score of all replicates using only data withheld for testing was high, with an AUC= 0.938. Standard deviations in AUC between each replicate run were minimal (lowest AUC = 0.937, highest AUC = 0.939). Additionally, environmental predictors deemed most important in explaining presence of Wilson’s Warblers were consistent across all replicate runs, with Bio 19 (Precipitation of the coldest quarter), Bio 5 (Maximum temperature of the warmest month), and Bio 4 (Temperature Seasonality) contributing to over 70% of the variation in occurrence explained by the MaxEnt model. Finally, our final map displaying the point-wise mean probability of occurrence (Fig. SX) closely matched Previously published maps of the species distribution of Wilson’s Warbler (for instance, probability of occurrence was consistently <0.23 in regions outside of the species range, and habitat suitability generally identified regions that were also found to harbor high abundance of Wilson’s warblers according to previously published data (Status of Birds in Canada, Peter Blancher, based on BBS abundance map estimates).

**Figure 1.**
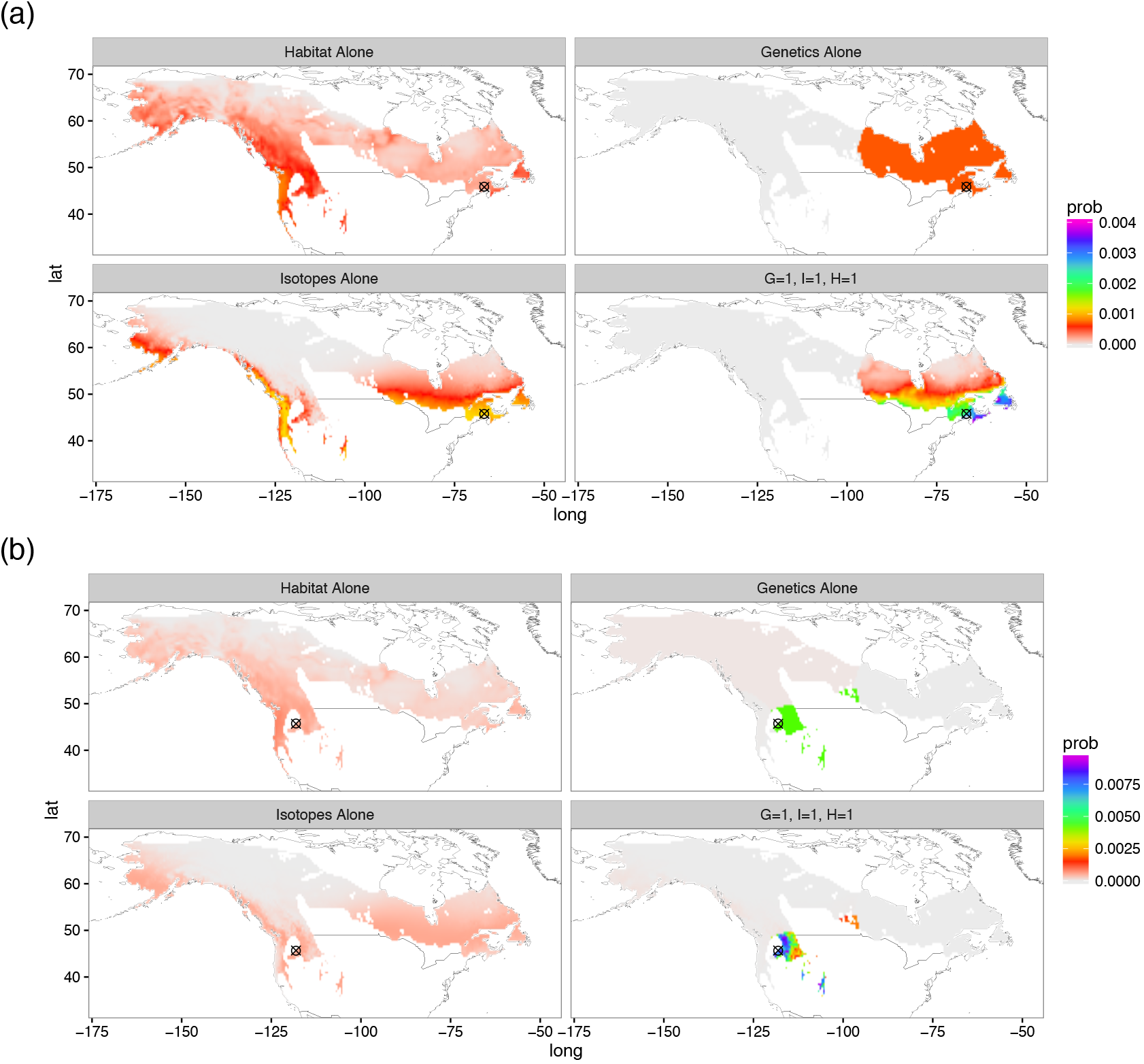
Representative posterior probability surfaces for 2 birds: (*a*) a bird from the Eastern region, and (*b*) a bird from the Rocky Mountains region. Each of the four panels shows the result using a different type of data. The first three ("Habitat alone," "Genetics Alone," and "Isotopes alone," show results for each data type applied separately. "*G* = 1, *I* = 1,*H* = 1" shows the results from the combined approach with *β*_iso_ = *β*_hab_ = 1. Maps which showed particularly strong examples of the combined approach are shown here, while maps for all 367 reference birds can be found in Supporting Information and to which we refer those interested in the variability in assignments across methods and individuals. In general, the maps support the idea that combining methods provides the most accurate assignment probabilities.

From predictor response curves, we found that increases in precipitation in the coldest quarter (Bio19) and temperature in the warmest month (Bio5) led to increases in the probability of Wilson’s warbler occurrence, whereas increases in temperature seasonality (Bio4) led to decreases in probability of occurrence (Fig. SX). These relationships resulted in several regions of high habitat suitability being identified within the Wilson’s warbler range, including much of the Pacific coastline, the Sierra Nevada Mountain range, parts of the Rocky Mountain range, and the Canadian Maritimes for eastern breeding populations.

### 3.4 Comparison of data types

Overall, combining genetics, isotopes, and habitat suitability improved the inference of breeding origin of Wilson’s warblers. The posterior mean great circle distance between the true location and inferred location for nearly every bird in the reference dataset was decreased by combining all three sources of data (Figure 2).

Of the three individual data sources, genetic data provided the most accurate localization of individuals, while isotope assignments and the habitat suitability prior used alone resulted in significantly less accurate localization. There were also significant differences in the accuracy of assignments of the reference birds to breeding location based upon geographic region (Figure 3): genetic assignments performed best in all regions; for birds originating from the Pacific Northwest, the Sierras and the Rocky Mountains, more accurate assignments were achieved by using the prior information from habitat suitability than using data on stable isotopes alone; and conversely isotope-only assignments outperformed assignment using just the habitat suitability prior in Coastal California and the Eastern United States.

### 3.d Example Data-Timing of migration in Pacific Northwest Wilson’s warblers

In order to illustrate our combined approach on real-world migratory data, we calculated the posterior mean remaining migratory distance of birds sampled from Cibola, AZ during the spring migrations of 2008 and 2009. A general pattern, previously suggested in Ruegg et al (2014), of birds en route to Coastal California migrating through before birds en route to the Pacific Northwest, the Sierras, and Alaska to Alberta, was reinforced using our combined-data approach (Figure 4*a*). In addition, we found previously undetected patterns in the timing of migrants en route to the Pacific Northwest, with migrants headed to the southern Pacific Northwest arriving earlier than migrants en route to northern Pacific Northwest. These results were concordant across both years (Figure 4*b*).

**Figure 2.**
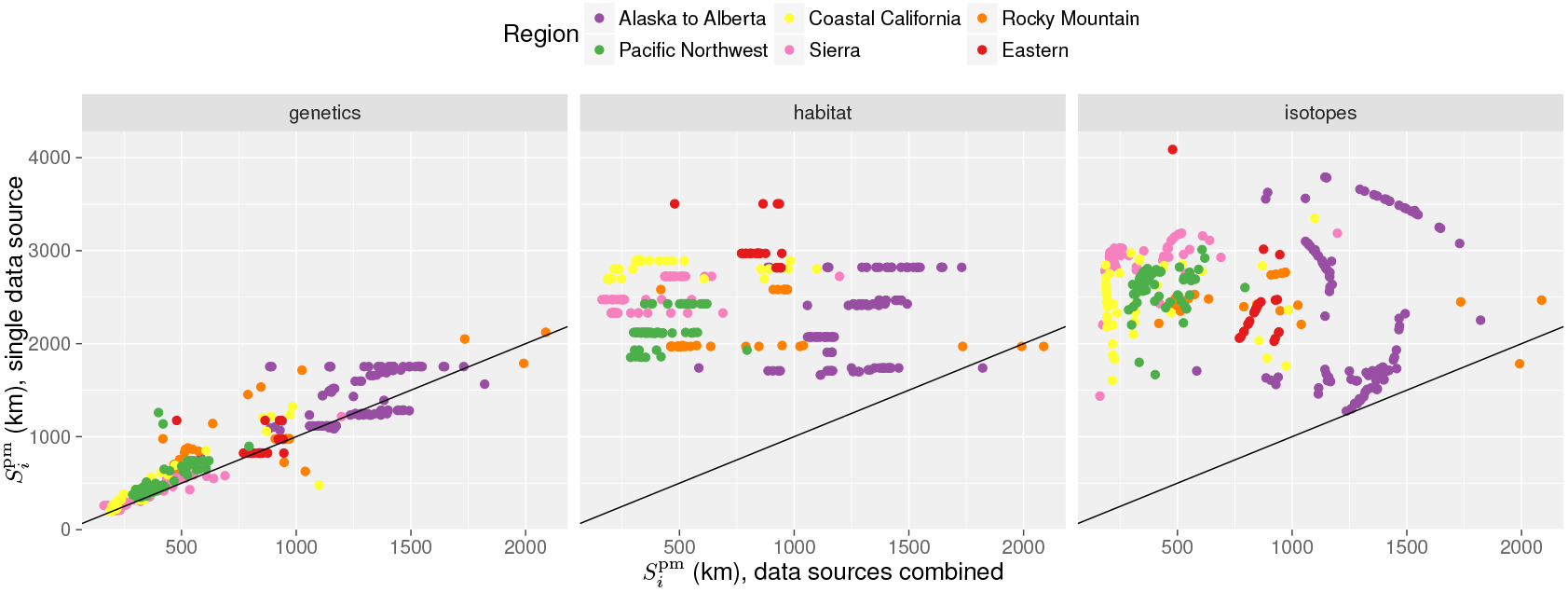
A comparison of the posterior mean great circle distances, 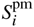, between individual data types and the combined approach. Each point represents one bird. In all panels the *x*-axis shows 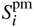 using all data sources combined and the y-axis shows 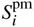 when using a single data source. The proximity of the points to the *y* = *x* line shows how well each method does compared to the combined approach. The results clearly demonstrate that genetic data provide the most accurate inference of the true origin, while habitat suitability and stable isotope data provide more modest improvements. Nonetheless the fact that the majority of points lie above the *y* = *x* line in the "genetics" plot confirms that the addition of habitat and stable isotope data improves the inference of breeding origin (as all the points would fall on the *y* = *x* line if the inclusion of stable isotopes and habitat did not change the inferences made with genetics alone).

**Figure 3.**
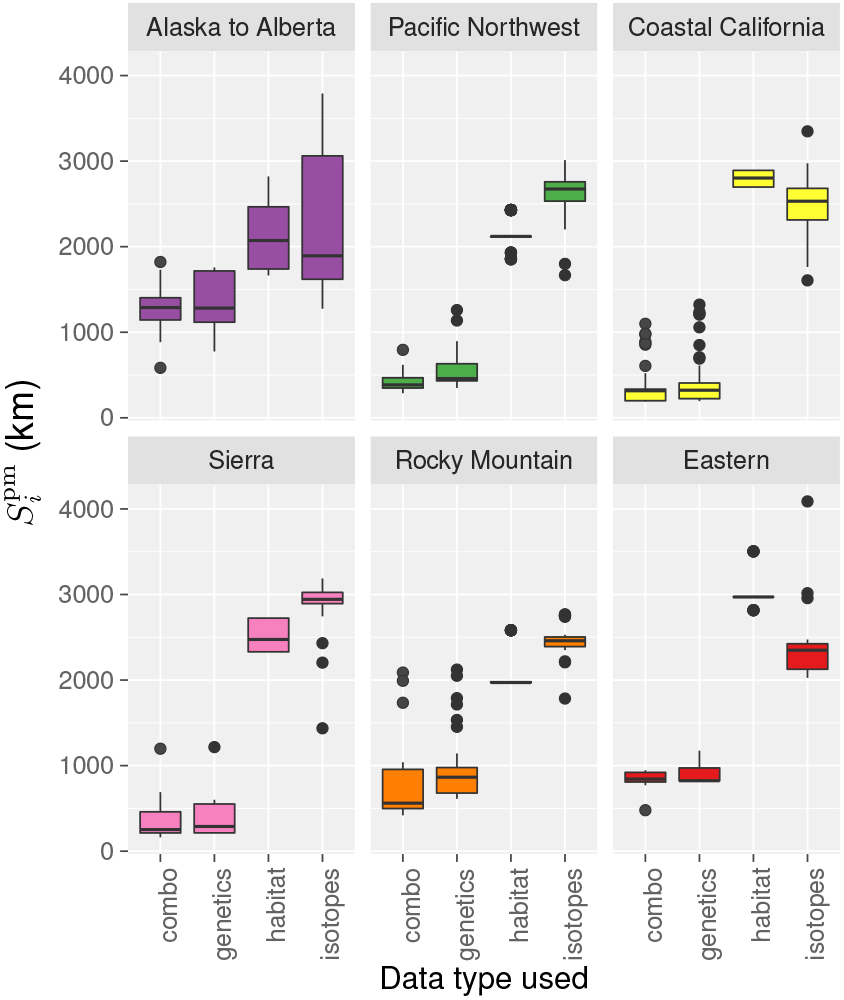
The accuracy of each type of data as estimated by the posterior mean great circle distance to the true location by region. The figure demonstrates that genetic data alone contributes the most to the accuracy of assignments to the true breeding location across all regions, while isotopic and habitat suitability data provide more modest improvements that differ in strength by region.

**Figure 4.**
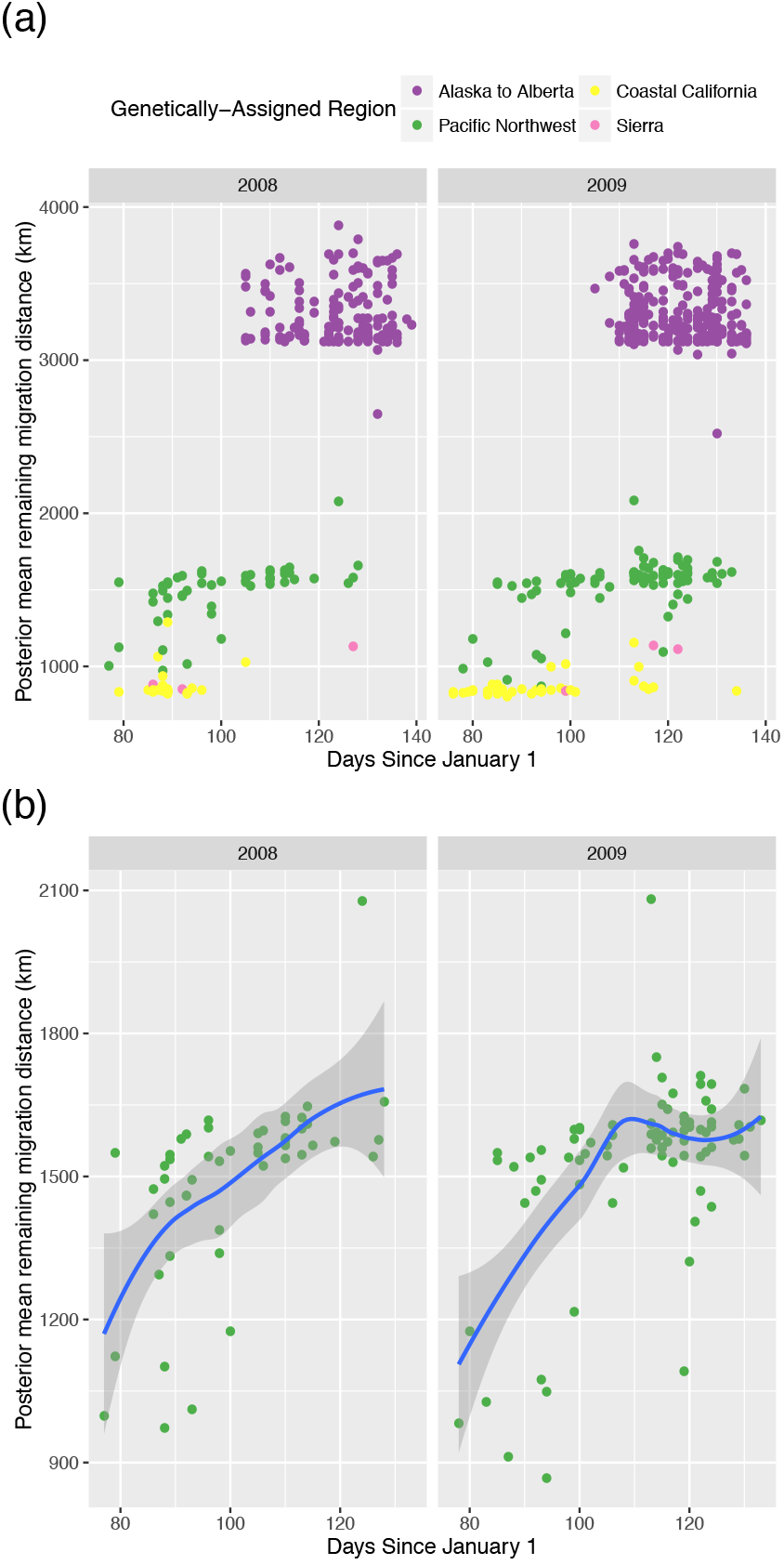
Utility of the combined approach demonstrated with spring Wilson’s warbler migrants stopping over in Cibola, AZ: (a) The posterior mean remaining migration distance by day of the year for all birds stopping over in Cibola AZ during spring migration across the years of 2008 and 2009. (b) The posterior mean remaining migration distance by day of the year for genetically-assigned Pacific Northwest birds stopping over in Cibola, AZ during spring migration, demonstrating a previously undetected pattern of birds en route to more northern regions of the Pacific Northwest arriving later than birds en route to the southern portion of the Pacific Northwest.

## 4. Discussion

Tracking the origins of migratory animals using non-invasive, intrinsic marking techniques has been a particularly challenging endeavor for movement ecologists. Here we develop a novel method for combining three independent data sources (genetics, stable isotopes, and habitat suitability models) that improves upon the accuracy of each method when used on its own. Using leave-one-out cross validation to compare the relative accuracy of each method independently, we find that genetic data alone provides the most accurate estimation of the true origin of our reference birds, consistently outperforming assignments based upon stable isotopes and habitat suitability models (Figure 1, Figure 3). Improvements to the R-code for batch isotopic and genetic assignments initially developed by Vander Zanden *et al*. (2014) and Anderson et al. (2008) respectively, were compiled into an R-package called gaiah in an effort to make future implementations of the resulting combined approach feasible across a broad range of migratory systems. Below we discuss region-and data-type-specific differences in the assignments, illustrate the utility of our combined method for uncovering new patterns of connectivity across time in Wilson’s warblers captured during spring migration, and discuss the implications of our results for future studies considering limitations in both time and resources.

### (1) Accuracy of each data type relative to the combined approach by region

A comparison between the posterior mean great circle distances using each method alone conclusively demonstrates that genetic-only assignments most closely approximate the true origin of birds in our reference sample (Figure 3, Figure 2). It is important to note that our analysis is based upon a single species and one might wonder whether our results are limited to Wilson’s warblers or whether they will be generalizable across other migratory systems. While the accumulation of additional data is needed to more thoroughly answer this question, results from numerous other molecular-based studies suggest that the increase in accuracy gained from using high-resolution genetic tags will have wide utility for population assignment in migratory animals far beyond our single species application (Nielsen *et al*., 2009; Hess *et al*., 2011; Nielsen *et al*., 2012; Ruegg *et al*., 2014). It is also important to note that in Wilson’s warblers the genetic variation was clustered into distinct regions, but in cases where genetic variation changes more gradually as a function of geographic distance (*i.e.*, isolation-by-distance), it would be possible to construct the genetic posterior matrices, 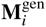, by using an assignment approach designed to deal with continuous spatial data (Wasser *et al*., 2004; Rañola *et al*., 2014). Such an approach is implemented in the package IsoScat (Rundel *et al*., 2013), but could be improved by incorporating a genetic model that allows for allele frequencies to change non-linearly with geography (*e.g.*, Rañola *et al*. 2014). Finally, the improvement in assignments based on genetic data alone are not surprising, considering that inferences based upon high-resolution genetic tags are coming from 96 axes that have been selected for informativeness in population level assignments, while inferences based upon stable isotopes are traditionally based upon only one or a few axes.

While high-resolution genetic markers alone provide a powerful framework for population-level assignment, the inclusion of stable isotope and habitat suitability data served to increase the precision and accuracy of breeding origin estimates in almost all cases (Figure 3). Surprisingly, for birds in some regions, such as the Rocky Mountains, the Pacific Northwest and the Sierras, simply using the habitat suitability prior alone provided a better inference of the true origin than did stable hydrogen isotope data. This is largely driven by the fact that continental variation in stable hydrogen isotopes in precipitation in North America is greater across latitudinal than longitudinal gradients (Hobson & Wassenaar, 2008), resulting in similar stable hydrogen isotope values between the Pacific Northwest and Sierra regions and comparable longitudes in the Rocky Mountains. Habitat suitability models have more power than stable isotopes in delineating among regions of similar longitudes that have high environmental heterogeneity, such as in the interior western regions of the United States. Whereas, in ecologically more homogeneous regions that vary across latitudes such as Alaska to Alberta, Coastal California, and Eastern North America, stable hydrogen isotope data provide more information about the true region of origin than does the inclusion of habitat suitability models.

Despite improvements in assignments using a combined approach, it still remains difficult to accurately identify the origin of individuals from breeding areas within the region from Alaska to Alberta using our methods. Genetic data alone provides only a very coarse inference of the geographic origin, with the spatial scale of assignments to this region being at best on the order of several thousand square kilometers. As a result, the data in Figure 4a appear discontinuous because our assignments are based upon the posterior mean remaining migration distance and the region from Alaska to Alberta is large. The inclusion of habitat suitability and stable isotope data helps little, mostly due to the fact that habitats are equally suitable and stable isotope values are similar in a large area of South-Central Alaska and Southern Interior British Columbia. Preliminary, unpublished results from the genetic data set reported in Ruegg *et al*. (2014) suggest that additional genetic differentiation may exist in this region and it is possible that increased genetic sampling could reduce the spatial scale of assignments. Alternatively, birds breeding from Alaska to Alberta may form a more homogeneous population in general, with birds having more flexibility in the timing of migration, in which case increased genetic sampling may provide limited benefits.

### (2) Demonstration of the utility of the combined approach

We demonstrated the utility of the combined method by assigning 686 Wilson’s warblers migrating through Cibola, AZ during the spring of 2008 and 2009, back to their most probable breeding destination. The results identify previously undetected patterns in the timing of migrants en route to the Pacific Northwest, with Wilson’s warblers en route to more southern locations in Northern California migrating through earlier than birds en route to more northern locations near the Washington, British Columbia boarder (Figure 4b). These results were consistent across years and further corroborate similar patterns of leap-frog migration in western Wilson’s warblers identified in previous work on a much coarser continental scale (Clegg *et al*., 2003; Paxton *et al*., 2007, 2013; Rundel *et al*., 2013). Interestingly, similar leapfrog patterns were not seen in migrants en route to breeding regions ranging from Alberta to Alaska (Figure 4), but is not clear whether this is due to a lack of resolution in the markers or the lack of a leap-frog pattern in this region. In addition, similar leapfrog patterns were also not observed within the California population where genetic resolution was high and stable isotopes provided the second most informative measure of the true location of reference birds. Overall these data illustrate the power of the combined approach for identifying fine scale patterns in migratory behavior which have not been detected using any other methodology to date.

### (3) Recommendations for future migratory-connectivity studies

Our analyses revealed that if operating under time or cost restraints, assignment accuracy using genetics alone can provide a nearly optimal estimation of accuracy without the need to add additional data sources. This is particularly true in cases where the volume of individuals to be screened is high (1000’s of individuals) because at that point the per-individual cost is less. Because of the ease with which high volumes of samples can be screened, intrinsic markers provide a powerful tool for statistically rigorous assessments of migration phenology. Despite this fact, it is important to note that the development of high-resolution genetic markers requires a substantial initial investment in highly-trained personnel able to process genome-wide data, as well as in sequencing and reagent costs. As such, this method may not be feasible in all situations. This initial investment is likely to pay dividends in the long-term, however, and the utility of SNPs for population assignment cannot be understated. Once a basemap of genetic variation across geographic space (i.e. a genoscape) has been produced, it can provide an extremely valuable resource for tracking populations and documenting likely range shifts due to climate change or other anthroprogenic stressors. In addition, the development of these markers can be utilized for a number of other purposes such as the identification of biologically meaningful population boundaries, studies of introgression, hybridization, parentage, kinship and effective population size among others (Andrews *et al*., 2016; Garner *et al*., 2016).

Despite the broad utility of genetic markers, many research projects will be faced with limitations in either time and/or resources. In these cases, our analyses indicate that a combination of habitat suitability and stable isotope data provides the second best option to genetics alone (SI 1, Fig. 2). Our results suggest that the only region where the inclusion of habitat suitability did not improve the accuracy of our assignments was in Eastern North America. In this region, the more the suitability models were weighted, the less accurate the resulting assignments became. We suspect a number of reasons for this trend; 1) records of our study species were much more limited in this region, leading to less accurate species distribution probabilities in eastern habitats, 2) a single species distribution model was created as a prior across the entire breeding range, which led to a model that was too general to capture nuances in the eastern part of the breeding range, particularly because a separate study using habitat suitability models for the Wilson’s warbler found little niche overlap between eastern and western groups, even suggesting that they might be considered cryptic species (Ruiz-Sánchez *et al*., 2015) and 3) eastern reference individuals were not sampled uniformly across the actual breeding range, leading to low-deviance (but inaccurate) estimates of origin (see Figure 3).

Even with these limitations, habitat suitability models based on individual occurrence records have proved to be one of the most effective tools for understanding species distributions, ecological requirements, and realized niches (Elith *et al*., 2011; Harrigan *et al*., 2014). As such we recommend their inclusion whenever possible. Habitat suitability models are also likely to be the easiest of the three described methods to apply for a given study species: their only required input is records of animal locations and a suite of environmental predictors for the study region. However, ease-of-use should be tempered with caution when utilizing these methods as priors. For instance, habitat suitability priors may purposefully down-weight regions that could represent actual origin locations of individuals, particularly when species are capable of surviving in marginal habitats, or where estimates of habitat suitability and population abundances do not necessarily align. It is vital that occurrence data be sampled adequately and in a manner representative of the entire habitat available to the species, as sampling effort and intensity can greatly affect final model results (as seen in our eastern individuals) (Phillips *et al*., 2006; Elith *et al*., 2011). Understanding the ecological mechanisms of the study species, and an informed interpretation of the results of any species distribution model is paramount, and can often lead to surprising insights (Elith *et al*., 2011; Renner & Warton, 2013; Harrigan *et al*., 2014). Despite these caveats, our method suggests that the inclusion of habitat suitability model output as a prior in assignment tests could prove valuable to increase the accuracy of estimates.

Well established patterns of stable hydrogen isotopes in precipitation that correspond to isotopic signatures in animal tissues (Cryan *et al*., 2004; Hobson & Wassenaar, 2008; Hobson *et al*., 2012b) have facilitated the widespread use of stable isotopes to understand migratory connectivity in birds (Hobson *et al*., 2010), bats (Cryan *et al*., 2014), butterflies (Hobson *et al*., 1999; Brattström *et al*., 2008), and dragonflies (Hobson *et al*., 2012a). Like genetic sampling, the collection of tissue samples (e.g., feathers, hair, insect chitin) for isotopic analysis does not require extensive training, and collection and analysis of a large numbers of samples is fairly inexpensive compared to extrinsic tracking devices. While it is important to understand the relationship between the tissue sampled and patterns of stable isotopes in the environment, technical expertise in stable isotope preparation and analysis is not necessary because of the accessibility of reputable isotope labs using a comparative equilibrium approach for the analysis of stable hydrogen isotope samples (Wassenaar & Hobson, 2003). Moreover, analytical tools such as ISoMAP (Bowen *et al*., 2014) and the R-package developed in this study increase the accessibility of technically challenging stable isotope analyses that incorporate error and prior probabilities to more scientists. These advantages have allowed for the coupling of stable isotope data with a wide range of disciplines, including neural biology (Barkan *et al*., 2016), endocrinology (Covino *et al*., 2016), and disease ecology (Gunnarsson *et al*., 2012; Ogden *et al*., 2015), greatly advancing our understanding of the ecology of migratory species throughout their annual cycle. Nevertheless, as demonstrated in this study the accuracy of stable isotopes alone to assign individuals to specific locations is much lower than when combined with high-resolution genetic markers and habitat suitability. The use of stable isotopes alone is most successful when assessing patterns of migratory connectivity at broader spatial scales (e.g., geographic regions), and for migratory species with geographic distributions that span large latitudinal gradients with little longitudinal variation (Hobson & Wassenaar, 2008). The combination of stable hydrogen isotopes with other intrinsic markers that have more longitudinal structure such as other stable isotopes (e.g., Sellick *et al*. 2009), morphometrics (Delingat *et al*., 2011), and genetic data (Clegg *et al*. 2003; Chabot *et al*. 2012, this study) can increase the accuracy of assignment. Additionally, longitudinal ambiguity can be reduced by constraining assignments to smaller geographic areas within a species overall range (e.g. western breeding locations only) based on known flyways or banding occurrence data that preclude assignment to some areas within a species range (Hobson *et al*., 2009; Van Wilgenburg & Hobson, 2011).

### (4) Conclusions

Identifying the population of origin of migratory animals using intrinsic-marker techniques is now feasible at increasingly small spatial scales. Here we show that genetic assignments far outweigh the accuracy of assignments based on isotopes or habitat suitability models alone. When logistically possible, the inclusion of all three data sources (genetics, stable isotopes, and habitat suitability) can, as in the case of the Wilson’s warbler, serve to refine genetic-only estimates and reveal previously undetected patterns in the timing of migratory events. Initial (seemingly large) investments in developing high-resolution genetic markers must be weighed on a case-by-case basis, but undoubtedly provide a high return on investment for heavily managed species or species for which high volumes of samples are to be screened. Our results provide methodological recommendations and a framework for analysis that can be used to facilitate future advances in the field of movement ecology including the discovery of new migratory pathways and cryptic migratory species, as well as the tools necessary to manage declining taxa in the face of rapid ecological changes.

## Acknowledgements

We thank Eli Bridge and Hanna Vander Zanden for discussions about isotopic assignment test methods and for sharing the R-code they used to perform isotope assignments and to Hanna Vander Zanden for her comments on an earlier version of this manuscript. We also thank the Intsitute for Bird Populations and MAPS, MoSI, and LaMNA networks for sample coordination and contribution. Funding for this work was made possible by grants from the California Energy Commission to K Ruegg (POEA01-Z01) and to K Ruegg and TB Smith (EPC-15-043) as well as grants from the National Science Foundation to R Harrigan (PD-08-1269) and F Moore (IOS-0844703). This work was also supported by a grant from the Environmental Protection Agency (R 833778) to TB Smith and generous donations from Margery Nicolson, an anonymous donor, and First Solar Inc.

## Appendix A Parametric bootstrap for calculation of isotope posterior probabilities

Here we describe the approach of Vander Zanden *et al*. (2014) for using the 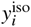 values of birds sampled from known breeding locations in a parametric bootstrap procedure to derive values of 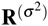, 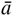, and 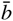. Let *ℓ* = 1,…,*L* index *L* different sampling locations from which at least two breeding birds were sampled for isotopes. Nearby locations can be merged into a single location if they are all in the same cell of the grid 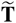. For each location *ℓ* we computed the mean and standard deviation of the 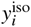 values of the breeding birds there, *i.e.*,

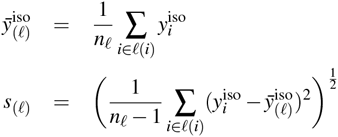

where *n_ℓ_* is the number of breeding birds sampled at *ℓ* and *ℓ*(*i*) is the set of their indices. Then, to create a parametric bootstrap sample of the isotope data from the reference birds, at each *ℓ*, *n_ℓ_* values of *y_i_* were simulated from a normal distribution with mean 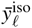 and standard deviation *s*_(*ℓ*)_. This bootstrap sample was then regressed as the dependent variable against the values in **P**(*^μ^*) corresponding to each location *ℓ*, as the independent variable, yielding, for each bootstrap replicate a linear relationship, parameterized by a slope and an intercept, between the predicted isotope ratios from IsoMap and those observed in birds from each location. The average over bootstrap replicates of the these slopes and intercepts, respectively, are used as 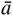 and 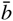 in Equation 1. The elements of the matrix 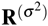 in (2) are the variances over the bootstrapped values of the predicted feather isotope values. That is, for each row *r* and column *c*:

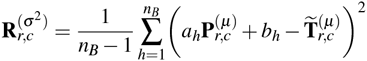

Where *a_h_* and *b_h_* are the slope and intercept estimated from the *h*^th^ of *n_B_* bootstrap samples.

